# Modified self-amplifying RNA mediates robust and prolonged gene expression in the mouse and ex vivo human brain

**DOI:** 10.1101/2025.10.30.685635

**Authors:** Jennifer Freire, Sicheng Pang, Joshua E. McGee, Maxwell J. Heinrich, Jiancheng Xie, Elijah Hammarlund, Dana Shaw, Yuxin Zhou, Yangyang Wang, Colin Porter, Lauren Dang, Erynne San Antonio, Ziqing Yu, Kexin Li, Scellig Stone, Hart Lidov, Jordan Farrell, Emily K. Osterweil, Wilson W. Wong, Mark W. Grinstaff, Xue Han

**Affiliations:** Department of Biomedical Engineering, Boston University, Boston, MA, USA; Department of Pharmacology, Physiology, and Biophysics, Boston, University, Boston, MA, USA; Rosamund Stone Zander and Hansjoerg Wyss Translational Neuroscience Center and F.M. Kirby Neurobiology Center, Boston Children’s Hospital, Boston, MA, USA; Graduate Program for Neuroscience, Boston University, Boston, MA, USA; Undergraduate Program for Neuroscience, Boston University, Boston, MA, USA; Department of Neurosurgery, Boston Children’s Hospital, Harvard Medical School, Boston, MA, USA; Department of Neurology, Boston Children’s Hospital, Harvard Medical School, Boston, MA, USA; Biological Design Center, Boston University, Boston, MA, USA; Department of Chemistry, Boston University, Boston, MA, USA

**Keywords:** Immune evasion, central nervous system, modified saRNA, gene transfer, astrocytes, neurons, human brain slice cultures

## Abstract

Facile, non-genomic integrating gene delivery technologies are lacking for rapid onset and prolonged protein expression in the brain. Here we report the protein expression and cell type tropism for an advanced messenger ribonucleic acid (mRNA) technology, modified 5-hydroxymethylcytidine (hm5C) self-amplifying ribonucleic acid (saRNA), when injected into the mouse brain or applied to *ex vivo* human cortical brain slices. saRNA, encoding fluorescent proteins, encapsulated in an LNP formulation comprising ALC-0315 (present in Comirnaty®) efficiently mediates long-lasting protein expression in mouse brain cells beyond five weeks, with detectable expression in some neurons at three months. hm5C saRNA substantially outperforms N1mΨ mRNA. In addition to transfecting astrocytes and neurons at the injection site, hm5C saRNA-LNPs label neurons retrogradely. Excitingly, hm5C saRNA-LNPs afford protein expression in human cortical brain slices, with expression emerging within 24 hours and lasting beyond 76 days. Modified saRNA provides new opportunities for mechanistic neuroscience research and therapeutic development.

## INTRODUCTION

The ability to genetically modify brain cells is advancing our understanding of the brain and offers exciting therapeutic possibilities (*1–4*). While viral transduction remains the most effective method for nucleic acid delivery in the central nervous system, it presents significant limitations in payload capacity and safety concerns for clinical applications. Nonviral ribonucleic acid (RNA) technologies offer promising potential to advance the therapeutic development for neurological and psychiatric diseases (*1, 5, 6*). Unlike deoxyribonucleic acid (DNA), messenger ribonucleic acid (mRNA) does not require nuclear entry, thereby eliminating the risk of genomic integration associated with DNA-based methods. The development and successful clinical outcomes of mRNA-based COVID vaccines, which incorporated the groundbreaking discovery of chemically modified nucleotides (N1mΨ) to reduce mRNA immunogenicity and enhance translation efficiency, highlight the potential of mRNA therapeutics (*7–10*).

Past efforts to deliver mRNA encapsulated in lipid nanoparticles (LNPs) to brain cells demonstrate technical feasibility, though with limited transfection efficiency and a short half-life. For example, a landmark study in 2001 by Kariko *et al.* reported that injecting mRNA encapsulated in cationic lipofectin into the rat brain led to immediate protein expression within 1 hour, but the expression became undetectable after about 24 hours (*11*). More recent attempts in delivering mRNA loaded LNPs to brain cells intracerebrally or intraventricularly report improved transfection efficiency (*12–16*), but nonetheless fail to mediate long lasting gene expression due to the inherent limitation of the short half-life of the mRNA transcripts.

Self-amplifying RNAs (saRNAs) encode an RNA-dependent RNA polymerase that drives intracellular self-replication, resulting in sustained production of protein encoding RNA (*17, 18*). Intracerebral administration of a heme oxygenase-1 encoding wild-type saRNA, encapsulated in deoxycholic acid-conjugated polyethylenimine, affords persistent protein expression for over 7 days in the brain (*19*). However, wild-type saRNA triggers a type I interferon response that degrades the RNA transcript, limiting its utility despite extensive exploration for vaccines and therapeutics (*20*). Past attempts to reduce saRNA immunogenicity, including incorporating N1mΨ modification, failed to preserve transfection potency (*17, 21–29*). Recent studies show that complete substitution of cytidine with 5-hydroxymethylcytidine (hm5C) or 5-methylcytidine (m5C) reduces immunogenicity and affords prolonged protein expression (*30, 31*). When encoded for the SARS-CoV spike protein and packaged in LNPs, hm5C saRNA expresses several folds more spike protein in muscle cells than unmodified saRNA and N1mΨ mRNA, while reducing inflammatory cytokine and interferon alpha production (*30*).

Motivated by the improved safety profile and transfection efficiency of intramuscular administration of hm5C modified saRNA, we evaluated if modified saRNA-LNPs transfect brain cells when delivered intracerebrally. We compared leading LNPs and identified a formulation comprising ALC-0315 (present in Comirnaty, BioNTech/Pfizer) as an effective LNP, and then systematically evaluated the time course and cell type tropism of these saRNA-LNP mediated fluorescent protein expression in the mouse brain. In addition to examining expression in mice, the therapeutic potential of this technology relies on effective application to the human brain. To demonstrate this translational potential, we additionally tested saRNA-LNPs in *ex vivo* human cortical brain tissue prepared from surgical resections from epilepsy patients.

## RESULTS

### ALC0315 containing LNPs effectively deliver saRNA to mammalian brain cells

To evaluate the transfection efficiency of saRNA in the mammalian brain, we first tested four leading LNP formulations, comprised of an ionizable lipid, DOPE, cholesterol, and DMG-PEG2K at molar ratios of 50:10:38.5:1.5(*30, 32, 33*). The ionizable lipids were SM102 (Spikevax®, Moderna), ALC-0315 (Comirnaty®, BioNTech/Pfizer), L319, and Lipid A9. We loaded hm5C modified saRNA encoding either mCherry or GFP into these LNP formulations to allow for parallel screening of two LNP formulations in the same mouse. The assembled saRNA-LNPs possess a size between 100-200nm (Fig. 1B), a polydispersity index <0.2 (Fig. 1C), and an encapsulation efficiency >80% (Fig. 1D). We intracerebrally injected these saRNA-LNPs into the striatum of mice (625 ng, 5µL at 125 ng/µL at three depth, Methods, Fig. 1E) to maximally label the brain regions around the injection site, and the mCherry and GFP encoding saRNA-LNPs were injected separately in the two hemispheres in each mouse. We then collected and sectioned the brains three days post injection. Three experimenters visually inspected mCherry or GFP expression across brain sections blinded to experimental conditions, and noted fluorophore expression mediated by A9, ALC0315, and L319, but not by SM102 based LNPs (Fig. 1F–H). The expression levels, measured as area positive for fluorophore signals in the striatum area, are 12.5% for ALC-0315, 7.3% for A9, and 5.5% for L319 (5.52%) (Fig. 1F, Hii). Thus, we selected LNPs containing ALC-0315 for further testing.

**Figure 1.**
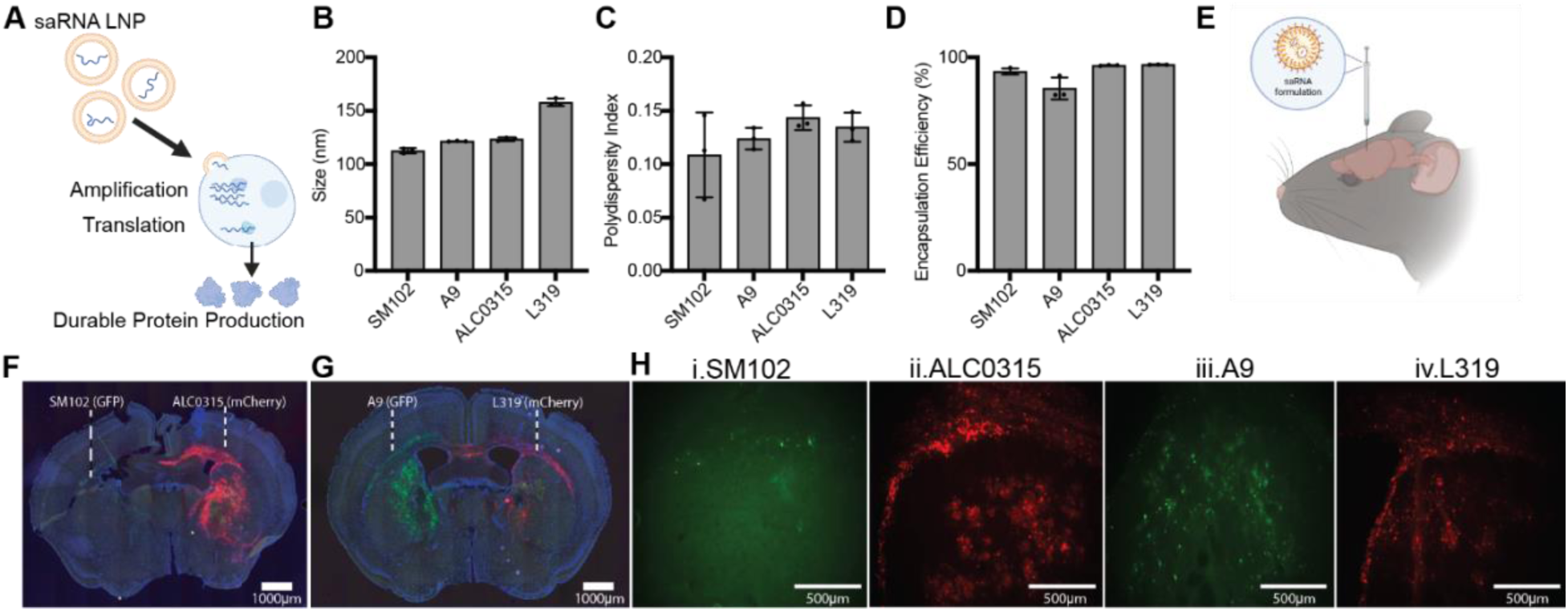
Testing LNP formulations for efficient saRNA delivery into brain cells. **(A)**Conceptual illustration of saRNA-LNP mediated protein expression. Created in BioRender. Pang, S. (https://BioRender.com/8hw84oi) is licensed under CC BY 4.0. **(B-D)** Size (**B**), polydispersity (**C**), and encapsulation efficiency (**D**) of LNP formulations comprising SM102, A9, ALC-0315, and L319 (mean±standard deviation, n=3 measurements). **(E)** Illustration of experimental testing in the mouse brain. Created in BioRender. Freire, J. (https://BioRender.com/inehs9l) is licensed under CC BY 4.0. **(F, G)** Fluorophore expression mediated by four LNPs loaded with saRNA encoding GFP or mCherry. Shown are GFP (green), mCherry (red), and DAPI (blue) fluorescence at 3 days post saRNA-LNP injection. **(H)** Zoom-in views of F and G showing GFP (green) and mCherry (red) fluorescence.

### saRNA-LNPs mediate prolonged gene expression in the mouse brain

To quantify the efficiency and the time course of saRNA-LNP mediated gene expression, we injected the ALC-0315 based LNP composition loaded with saRNA encoding mCherry into the mouse striatum unilaterally at three depths as in Fig. 1E (total 5µL at 125 ng/µL) to maximally label the brain area around the injection site (n=21 mice total). As a comparison, another cohort of animals received ALC-0315 LNPs loaded with conventional N1mΨ mRNA encoding mCherry (mRNA-LNPs) at the same dose and locations (n=20 mice total). We collected and sectioned these brains after varying durations post injection, from 24 hours to 3 months, and examined mCherry fluorescence without any antibody staining on brain sections. One day after saRNA-LNP or mRNA-LNP injection, mCherry expression is present around the injection site and corpus callosum and persists for at least 2-3 weeks (Fig. 2A).

**Figure 2.**
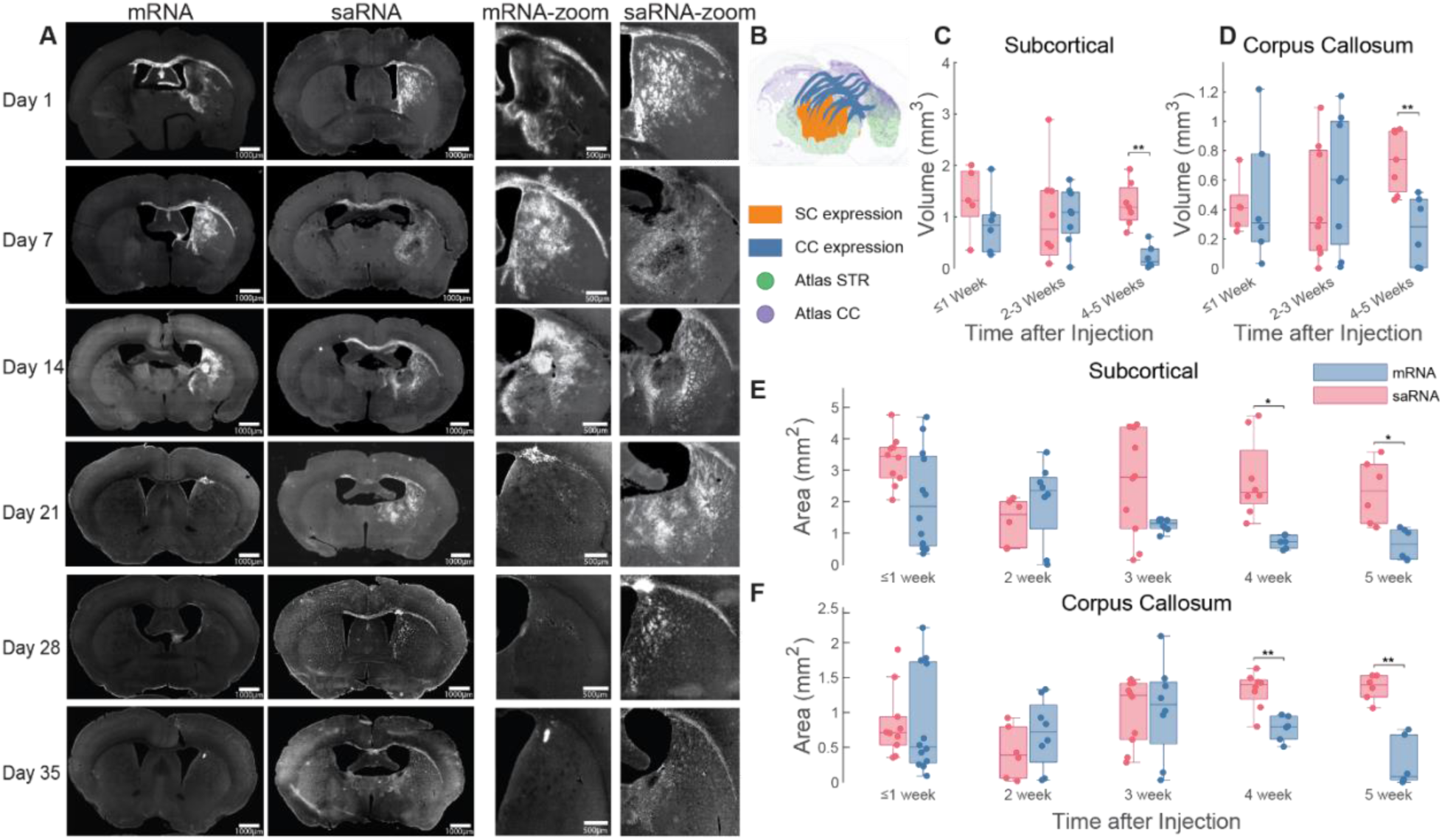
saRNA-LNPs mediate long-lasting gene expression. **(A)** Example brain sections at days 1–35 post-injection, showing mCherry fluorescence around the injection site in the striatum and throughout the corpus callosum. Left, whole-brain sections (scale bars, 1000 µm), with the injection into the right hemisphere for mRNA-LNPs or saRNA-LNPs; right, corresponding zoomed-in views around the injected area (scale bars, 500 µm). **(B)** 3D visualization of segmented mCherry-positive brain sections registered to the Allen Mouse Brain Atlas. SC: mCherry-positive subcortical region; CC: mCherry-positive corpus callosum region; Atlas STR: striatum in the Atlas; Atlas CC: corpus callosum in the Atlas. **(C, D)** Tissue volume positive for mCherry expression in the subcortical **(C)** and corpus callosum regions **(D)** across weeks after injection. n = 5–8 mice per time point per brain region. Significance was assessed with two-sided Mann–Whitney U tests. Statistical details in Table S1. **(E, F)** Peak area positive for mCherry in the subcortical **(E)** and corpus callosum regions **(F)** at varying time intervals post-injection; n = 2 slices per mouse, n = 3–6 mice per time point. Significance was assessed using a linear mixed-effects model with mouse as a random effect. All plots are box plots (box: 1st and 3rd quartiles; horizontal line: median; whiskers: 1.5× interquartile range). Statistical details in Table S2. ***, p < 0.005; **, p < 0.01; *, p < 0.05.

Because of the distinct morphology of subcortical regions versus the corpus callosum, we manually segmented the mCherry-positive area in the subcortical region and the corpus callosum separately on each brain section around the injection site. We then registered the segmented brain sections to the Allen Mouse Brain Common Coordinate Framework (Fig. 2B), and estimated the expression volume by interpolation and integration (Fig. 2B, C, Videos S1, S2). Within the first week and at two-to-three weeks, mRNA-LNPs and saRNA-LNPs mediate similar expression in the subcortical region around the injection site (≤1 week: 0.87 vs. 1.35 mm³, p = 0.18; 2–3 weeks: 1.03 vs. 1.00 mm³, p = 0.72, two-sided Mann-Whitney U test, statistical details in Table S1), and the corpus callosum region (≤1 week: 0.47 vs. 0.42 mm³, p = 0.79; 2–3 weeks: 0.59 vs. 0.45 mm³, p = 0.57). However, at four-to-five weeks, saRNA-LNPs afford significantly higher expression than mRNA-LNPs in both the subcortical (1.25 vs. 0.22 mm³, p = 0.0012), and the corpus callosum regions (0.73 vs. 0.26 mm³, p = 0.008, statistical details in Table S2).

To pinpoint when mCherry expression mediated by saRNA-LNP diverges from that by mRNA-LNP, we examined two brain sections with the highest expression from each mouse at a finer time duration post injection (Fig. 2D, E). mCherry expression mediated by saRNA-LNPs is significantly greater than mRNA-LNP as early as week 4 in both subcortical and corpus callosum regions (subcortical area, p=0.021; corpus callosum, p = 0.007, linear mixed-effect model, statistical details in Table S2). This protein expression divergence continues through week 5 (subcortical area, p = 0.024; corpus callosum, p = 0.001).

Finally, in one animal examined at three months post saRNA-LNP injection (the longest time-point evaluated), mCherry fluorescence remains present in some neurons, with strong labelling throughout the soma and neuronal processes (Fig. S1). The persistent saRNA-mediated transcript expression in neurons, though sparse, at three months after a single injection further underscores the effectiveness of saRNA mediated neuronal transfection. Together, these results demonstrate that hm5C modified saRNA mediates prolonged fluorophore expression at high levels, detectable with fluorescence microscopy without the need for antibody staining. saRNA mediates greater gene expression than mRNA after 4 weeks post injection and the expression remains strong at 5 weeks, while mRNA mediated expression starts to decay after 3 weeks.

### saRNA-LNPs transfect both neurons and glia, but with a preferential transfection of astrocytes at the site of injection

To determine the cell types transfected by saRNA-LNPs, we performed immunostaining using antibodies against the astrocyte marker protein GFAP and the neuronal marker protein NeuN (Fig. 3A-C). As astrocytes do not exhibit a well-defined morphology, we calculated the fraction of mCherry positive (mCherry+) pixels that are also GFAP positive (GFAP+). 43-56% of mCherry+ pixels are GFAP+ across the post injection time points examined. As neurons possess a defined morphology, we computed the fraction of mCherry+ pixels that are part of a neuron (NeuN+). Only ∼1-4% of labeled pixels are neurons across the time points examined, significantly lower than astrocytes (Fig. 3D). saRNA-LNPs transfect both neurons and astrocytes, with the majority of labeled cells being astrocytes at the striatal site of injection.

**Figure 3.**
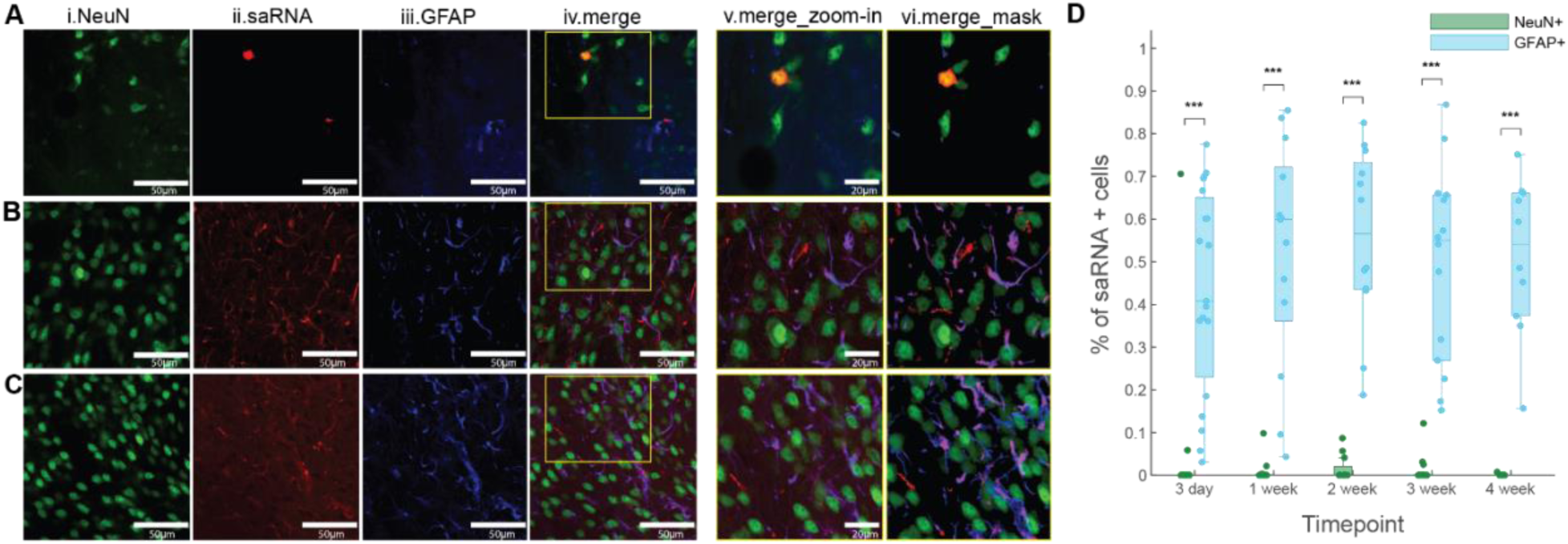
saRNA-LNP transfected more astrocytes than neurons in the striatum. **(A)** An example brain section showing saRNA-LNP mediated mCherry expression in neurons. (i) NeuN immunofluorescence from neurons (green); (ii) saRNA-LNP mediated mCherry fluorescence; (iii) GFAP immunofluorescence from astrocytes (blue); (iv) Merge, showing colocalization of mCherry and NeuN immunofluorescence (orange). Scale bar for (i-iv), 50 µm. (v). zoomed in view of the area highlighted by the yellow box in iv. Scale bars, 20 µm. (vi). Cell profiler identification of pixels positive for NeuN (green), mCherry (red), and GFAP (blue). **(B, C)** Similar to **A**, but for example brain sections showing saRNA-LNP mediated mCherry expression in astrocytes. Merge showing colocalization of mCherry and GFAP immunofluorescence (purple). **(D)** Quantification of the percentage of saRNA-mediated mCherry expression colocalization with neurons (green) versus glia (blue) across 5 time points after saRNA injection. ***, p<0.001, Wilcoxon rank-sum test (n=10-19 fields of view per timepoint, statistical details in Table S3).

### saRNA-LNPs mediate retrograde labeling of cortical neurons projecting to the striatal injection site

In addition to the expression at the site of injection, mCherry+ neurons are prominent in the cortex, both ipsilateral and contralateral to the inject site, starting about two weeks post injection (Fig 4A-C). Many labeled neurons exhibit bright mCherry fluorescence at the cell bodies, and throughout proximal and distal processes (Fig 4C). As broad cortical regions on both hemispheres, particularly on the contralateral side, project to the striatum, this observation suggests that saRNA-LNPs retrogradely label neurons.

**Figure 4.**
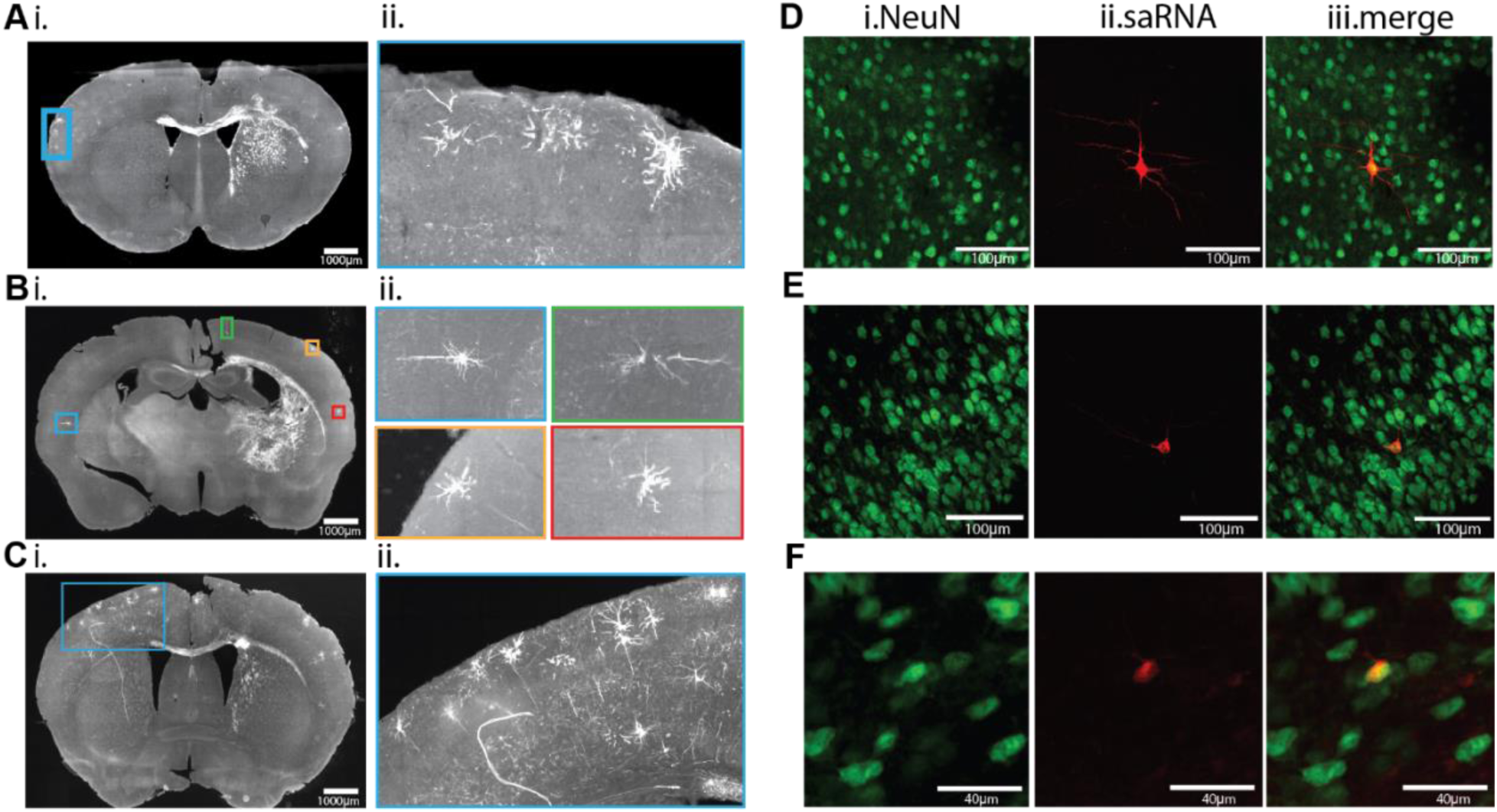
saRNA-LNPs retrogradely label cortical neurons. **(A)**(i) An example brain section at 2 weeks post injection showing mCherry positive cortical neurons contralateral to the injected striatum. (ii) A zoomed-in view of labeled neurons highlighted by the blue box in A. **(B)** An example brain section at 3 weeks post injection showing mCherry positive cortical neurons both contralateral and ipsilateral to the injected striatum. (ii) Zoomed-in views of labeled neurons across different areas of the two hemispheres highlighted in blue, green, yellow, and red boxes in **B**. **(C)**(i) An example brain section at 4 weeks post injection showing many mCherry positive cortical neurons contralateral to the injected striatum; same brain section image as shown in Fig. 2A. (ii) Zoomed-in view of labeled neurons highlighted by blue box in **C**. **(D-F)** Example fields of view showing saRNA-LNP mediated mCherry expression in cortical neurons. (i) NeuN immunofluorescence from neurons (green); (ii) saRNA-LNP mediated mCherry fluorescence (red); (iii) Merge, showing colocalization of mCherry and NeuN immunofluorescence (yellow).

### saRNA-LNP and mRNA-LNP lead to similar increases in astrocyte reactivity

To assess neuroinflammation associated with intracerebral administration of saRNA-LNPs and mRNA-LNPs, we examined astrocyte reactivity using antibodies against astrocyte marker protein GFAP. Since each mouse only received injections in one hemisphere, we compared between injected side versus the contralateral un-injected side. Specifically, we computed the fraction of pixels positive for GFAP ipsilateral versus contralateral to the injection site in 2-4 sections per animal at 2-3 weeks post injection. As a control, we injected PBS unilaterally (3 depth, and a total of 5uL) in a separate cohort of 3 animals and assessed GFAP immunofluorescence at 3 weeks post injection.

Across all groups, GFAP immunofluorescence is greatest around the injection site compared to the contralateral side (Fig. 5A, B). To quantify this, we computed the ratio of GFAP positive pixels ipsilateral over contralateral to the injection site, and confirmed that all three groups exhibit significantly greater GFAP expression on the ipsilateral side (each group vs. a ratio of 1, P < 0.05 for all groups, Wilcoxon signed-rank test; Fig. 5C, detailed statistics in Table S4), consistent with the general gliosis response to tissue injury from the injection procedure (*34, 35*).

**Figure 5.**
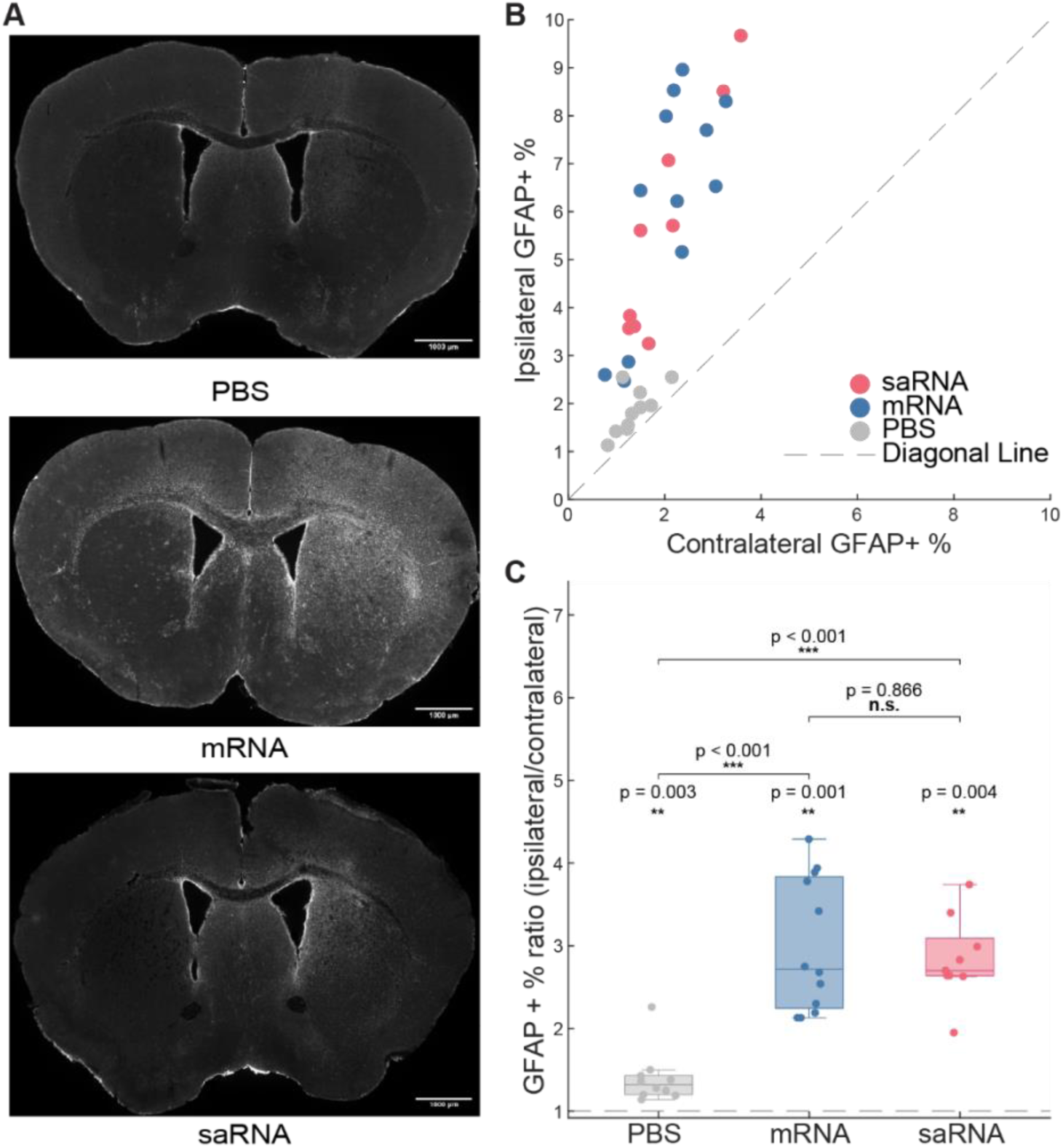
mRNA-LNP and saRNA-LNP intracerebral delivery increase astrocyte reactivity. **(A)** Example brain sections around the site of injection with PBS (top), mRNA-LNP (middle) and saRNA-LNP (bottom). Injection site is on the right. **(B)** Scatter plot of the fraction of GFAP positive pixels ipsilateral versus contralateral to the injection site. Each dot is one section (n=2-4 sections from 3-4 mice per experimental group). **(C)** Ratio of GFAP positive pixels ipsilateral over contralateral to the injection site. Each dot is one brain section (n=9-12 sections from 3-4 mice). For each box plot, the center line is the median, boxes are the interquartile range (IQR), and whiskers are 1.5× the IQR. The dashed line marks a ratio of 1. Each group’s difference from a ratio of 1 (stars above boxes) was tested with a two-sided Wilcoxon signed-rank test. Between groups were tested with linear mixed-effects model followed by Benjamini–Hochberg post-hoc correction. Statistical details are in Table S4. *P < 0.05, **P < 0.01, ***P < 0.001; n.s., not significant.

Comparing between mRNA-LNP, saRNA-LNP, and PBS treated groups, GFAP immunofluorescence is significantly higher in the mRNA-LNP and saRNA-LNP treated groups than the PBS treated group (mRNA-LNP vs PBS, p < 0.001; saRNA-LNP vs PBS, p< 0.001, linear mixed-effects model), and there is no difference between mRNA-LNP and saRNA-LNP treated groups (p = 0.866). In sum, saRNA-LNP and mRNA-LNP induce similar neuroinflammation at 2-3 weeks post injection that is beyond what is expected from injection induced gliosis.

### saRNA-LNPs mediate long lasting transgene expression in ex vivo human cortical brain slices

To explore the translational potential of saRNA, we applied 2-4µL of saRNA-LNP on the surface of each *ex vivo* human cortical brain slice (n=11 slices from 6 patients, age 3 month-21 years old, patient details in Fig. S3). We then visualized mCherry fluorescence at different time points after application and repeatedly in some slices while minimizing the impact of imaging on slice health and reducing the risk of contamination. mCherry expression occurs as early as 24 hours post saRNA-LNP application and persisted for many days (Fig. 6 and Fig. S3). Remarkably, a longitudinal assessment in one frontal cortex slice reveals consistently strong mCherry expression from 24 hours to 76 days post application (Fig. 6). Almost all examined mCherry positive cells, across all fields of view in all slices, exhibit neuronal morphology, suggesting a neuronal tropism.

**Figure 6.**
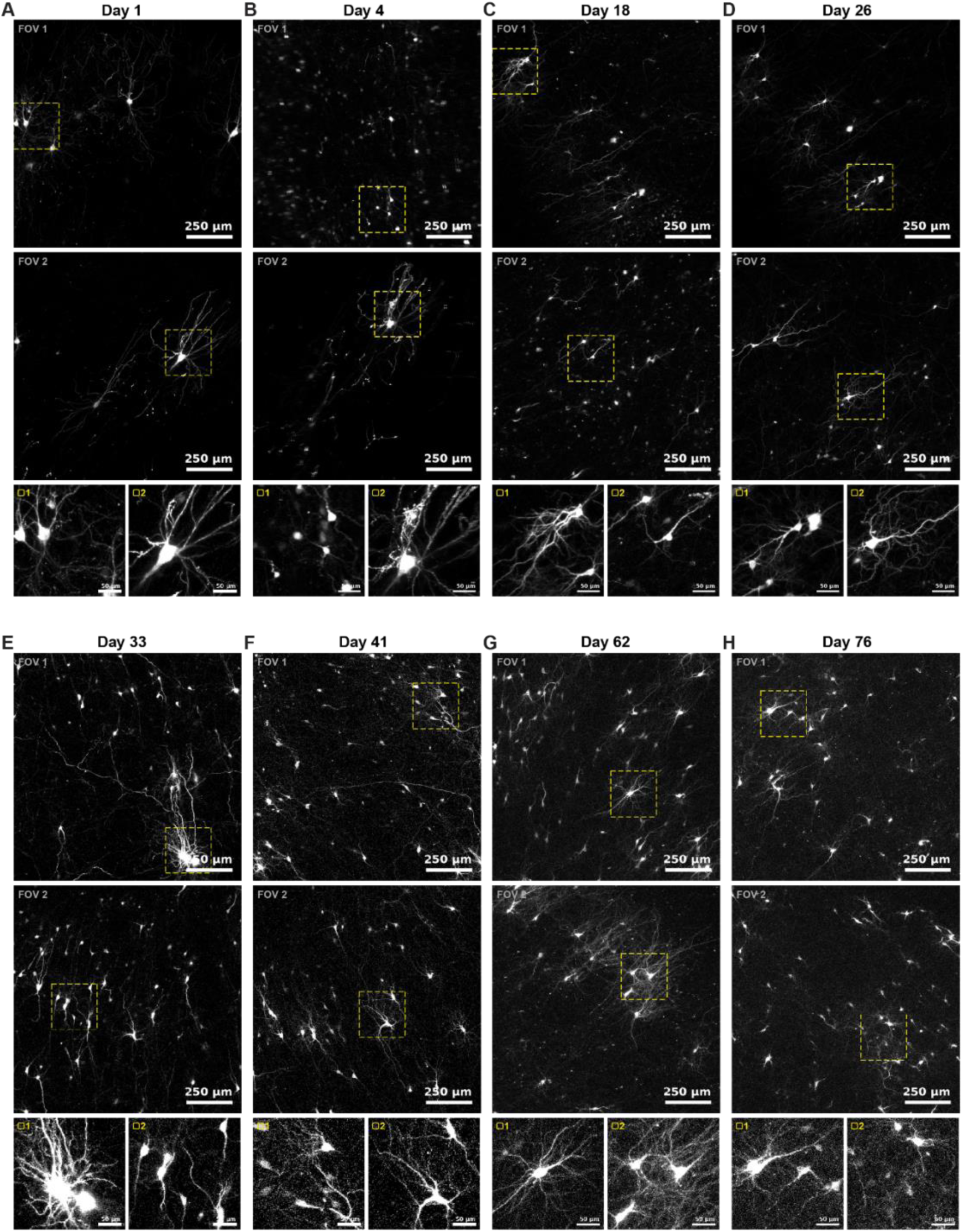
saRNA-LNPs mediate prolonged gene expression in ex vivo human brain slices. (**A-H**) Example fields of view showing mCherry fluorescence at 1-76 days post saRNA-LNP application in a temporal lobe brain slice obtained from 3-month-old male patient (Patient 1). Top two rows, two example fields of view (FOV1, FOV2; scale bars, 250 µm); bottom row, zoomed-in views of the region indicated by yellow dashed boxes in FOV1 (left) and FOV2 (right) (scale bars, 50 µm).

Because surgically resected tissue is inherently heterogeneous, varying in brain region, patient age, and slice health, we estimated transfection efficiency by computing the fraction of mCherry positive pixels per field of view. Across all fields of view examined, mCherry labelled pixels range from 0.36–2.41% (median 1.15%, IQR 0.94–1.63%, n=21 fields of view in 6 patients). For the one slice that we repeated imaged from 24 hours to 76 days post application (Fig. 6), the fraction remains around 1–2% across all time points assessed (Fig. S4, median 1.39%, IQR 1.44-1.66%, n=12 fields of view analyzed at 8 time points). Finally, we also applied mRNA-LNPs to one brain slice and detected mCherry expression at day 7 post application (Fig. S5). saRNA drives prolonged, neuron-tropic transgene expression in *ex vivo* human cortical slices despite substantial biological variability across slices, with a rapid expression onset within 24 hours of application and sustained expression beyond 76 days, highlighting the exciting translational potential of saRNA-LNP mediated gene delivery.

## DISCUSSION

Motivated by the durable expression previously observed from modified saRNA-based vaccine technology, (*30, 31*) we examined the transfection efficiency of hm5C saRNA encapsulated in ALC-0315 based LNPs following intracerebral delivery in the mouse and direct application on the surface of *ex vivo* human brain slices. In the mouse brain, saRNA-LNPs potently transfect brain cells at the injection site in the striatum region, and across broad cortical areas that project to the striatal injection site. Modified saRNA-LNPs afford prolonged mCherry transcript expression beyond five weeks, with some neurons remaining mCherry positive at three months post-delivery. hm5C saRNA yields significantly longer lasting protein expression than conventional N1mΨ mRNAs. Intriguingly, while hm5C saRNA-LNPs predominantly label astrocytes at the site of injection, many cortical neurons are also retrogradely labeled. In *ex vivo* human brain slices, prepared from samples obtained during standard surgical procedures, saRNA expression emerges within twenty-four hours and lasts beyond 76 days.

Modified hm5C saRNA expresses mCherry protein for over five weeks after a single injection, with mCherry fluorescence remaining prominent in some neurons near the injection site even at three months post injection. This is well beyond the expression mediated by mRNA previously reported (*11–16*), or the seven days observed with wild-type saRNA (*19*). Importantly, we directly measured mCherry expression without any amplification via antibody staining or enzymic reaction. This is in sharp contrast to previous studies that reported a Cre-mediated recombination process triggered by mRNA-mediated expression of Cre transcripts (*14, 15, 36*), which amplifies the transient presence of Cre expression into a more permeant recombination readout for easy of detection.

An important step toward translation is verifying that saRNA-LNPs drive effective expression in human brain tissue. To our knowledge, few studies have tested saRNA-LNPs in this setting, and transgene delivery in resected human tissue relies predominantly on viral vectors such as AAV. For eventual clinical application, saRNA-LNPs offer a more favorable safety profile than AAVs, eliminating the use of AAV capsid and the risk of insertional mutagenesis evoking cellular immune response (*37*). The genetic packing ceiling of saRNA is ∼ 15-20 kb, compared ∼ 5 kb for AAV, opening possibilities to deliver large proteins or multiple proteins (*3, 38*). Furthermore, saRNA-mediated expression onset is much faster than AAV, which requires days to reach peak transgene levels (*39*). This allows experiments to be performed early in the culture period, while tissue architecture and physiology remain closest to the native state, and makes it feasible to deliver genetically encoded calcium or voltage sensors and record from human neurons within a viable experimental window. Finally, the facile plug and play nature of saRNA to rapidly recode the sequence for expression of other proteins or multiple proteins is an advantage over viral vectors. Future work is needed to systematically compare the performance of saRNA and mRNA encapsulated in different LNPs, and to assess how donor age and underlying pathology affect saRNA-mediated expression in human brain tissue.

The retrograde labeling of cortical neurons that project to the striatal injection site, including from both the ipsilateral and the contralateral hemispheres, reflect axonal uptake of saRNA-LNPs. RNA-loaded LNPs will label neurons retrogradely *in vitro* (*40*), and the saRNA used here is derived from an encephalitis virus known to spread both anterogradely and retrogradely along axons (*41*).

Following axonal uptake, asRNA would need to be transported retrogradely to the soma, which may account for the slow emergence of mCherry-positive neuronal soma in cortical areas projecting to the striatal injection site. The fact that many labelled cortical neurons are in contralateral hemisphere, the intact un-injected side, rules out the possibility of direct transfection due to RNA-LNP leakage along the injection track. The same retrograde transport may also contribute to the mCherry expression observed in the corpus callosum, although RNA-LNP may directly transfect myelin, and future studies are needed to pinpoint the origin of corpus callosum labeling.

It is intriguing that hm5C saRNA-LNPs appear to label different brain cell types, predominantly glia cells in the subcortical mouse brain but neurons in human cortical slices. As we also observed retrograde labeling of cortical neurons projecting to the striatal injection site, it is possible that axon-mediated entry dominates in the human slice preparation, resulting in preferential labeling of neurons. In the mouse striatum, saRNA-LNPs may gain direct entry across the plasma membrane at the injection site. In addition, as intracerebral injection produces a local upregulation of reactive astrocytes, which may also contribute to increased glia labeling.

Around the striatal injection site in mice, the hm5C saRNA-LNPs predominantly label astrocytes and some neurons. Intracerebral injection of mRNA encoding Cre labels both astrocyte and neurons in the mouse striatum, as reported by Tuma et al (*36*). As the Cre-dependent recombination assay captures transient Cre-enzyme expression, the discrepancy in transgene expression in neurons may reflect the time course or the magnitude of variations between our study and Tuma *et al*. Indeed, the retrograde labeling of cortical neurons appears to progressively become stronger over the first couple of weeks, suggesting that saRNA transcripts may accumulate slower in neurons than in astrocytes, leading to the observed low protein expression in neurons at the site of injection. However, it is also possible that ALC-0315 LNPs exhibit a low transfection rate for the medium spiny neurons that dominate the striatal neuron population. Lastly, similar to the low microglial labeling reported previously for mRNA (*36, 42*), hm5C saRNA-LNPs do not readily transfect microglia as we only see limited instances of mCherry positive microglia (Fig. S2).

Chemically modified saRNA achieves durable expression by minimizing innate immune activation associated with unmodified RNA. Both RNA and ionizable lipids used to formulate RNA-LNPs are known to trigger systemic inflammatory responses (*43*), and the brain is particularly sensitive to injuries (*35, 44*). Indeed, we detected heightened astrocytes reactivity in the hemisphere injected with PBS control, mRNA-LNPs and saRNA-LNPs at two and three weeks post-injection, with mRNA-LNP and saRNA-LNP evoking stronger response than PBS control. Interestingly, hm5C saRNA-LNPs led to a similar level of gliosis at 2-3 weeks post administration as N1mΨ mRNA-LNPs, despite the substantially longer duration of mCherry expression mediated, indicating that the prolonged transgene expression afforded by hm5C saRNA does not incur greater astrocyte reactivity than standard mRNA. Current analysis is limited to astrocyte reactivity at two to three weeks post administration, future studies are needed to provide a more complete understanding of the evoked immunogenicity by assessing extended time points after injection and cellular response, which would elucidate the different transfection efficiency mediated by saRNA versus mRNA.

Recent advance of ionizable LNP designs yields improved mRNA delivery, particularly in vaccine development, offering promising potential for optimizing LNP-based non-viral gene transfection strategies for epithelial, connective, or neuronal tissues (*1, 6, 36, 45–48*). Ionizable lipids play a critical role in determining the physicochemical properties and delivery performance of the formulated LNPs. We tested four leading LNP formulations containing different ionizable lipids known to exhibit excellent transfection efficiency in human and mouse cells (*32, 33*). While formulations of SM102 (Spikevax®, Moderna) and ALC-0315 (Comirnaty®, BioNTech/Pfizer) effectively transfect muscle cells when formulated as vaccines (*33, 49*), their transfection efficiency in the brain varies, with SM102 transfecting fewer cells and ALC-0315 transfecting more. Future systematic testing of LNP compositions, such as using color compatible fluorophores, like GFP and mCherry demonstrated here, will help identify the most effective compositions for transfecting different brain areas or cell types. Aside from engineering LNPs to enhance mRNA delivery, incorporation of 3’UTR gene regulatory elements in the RNA sequence to bias gene expression between astrocytes and neurons can further control translation (*50*). Continued efforts on engineering LNP formulations and RNA sequence designs will further improve transfection efficiency and brain targeting specificity.

hm5C saRNA’s ability to mediate long-term gene expression in brain cells, in both mouse and human brain slices, presents profound therapeutic potential, e.g., to deliver glial derived neurotrophic factors for Parkinson’s disease (*51*), brain-derived neurotrophic factor for Alzheimer’s disease (*50*), Bcl-2 for ischemic stroke (*2, 52*), apoptosis-inducing ligand for glioblastoma (*53*), and thrombomodulin to boost blood brain barrier integrity(*54*). Additionally, continued progress to improve nonviral delivery of antisense RNA, siRNA, or CRISPR-Cas9 gene editing components into the brain will advance the ability to reduce risk factors of many neurological and psychiatric disorders (*1*), e.g., beta-secretase 1 or amyloid beta for Alzheimer’s disease (*55–58*), alpha-synuclein for Parkinson’s disease (*59, 60*), SOD1 for Amyotrophic lateral sclerosis (*61*), or as an immunotherapy for glioblastoma (*62, 63*). Thus, hm5C saRNA-LNP represents a potent non-viral gene transduction platform for the mammalian brain with broad applicability in basic neuroscience research and exciting translational potential.

## METHODS

### saRNA & mRNA LNP synthesis

Both saRNA and mRNA (GenBank accession number to be determined. Sequences will be deposited in GenBank upon manuscript acceptance) encoding mCherry or EGFP were synthesized by *in vitro* transcription with the T7 High Yield kit (New England Biolabs). For the saRNA constructs, cytidine was replaced by 5-hydroxymethyl (TriLink). For the mRNA constructs, uridine was replaced by N1-methylpseudouridine. Following transcription for 3 hours at 37 °C, the DNA templates were digested by TurboDNase (Invitrogen). The RNA preparations were purified using the Monarch RNA purification kit (New England Biolabs) and eluted in water. Additionally, cellulose purification was performed as previously described to remove double-stranded RNA contaminants (*64*).

The hm5C saRNA and N1mΨ mRNA encoding mCherry or EGFP were encapsulated into LNP formulations using ionizable lipids including ALC-0315 (Avanti), SM102 (Cayman), Lipid A9 (Cayman), or L319 (Cayman), in combination with DOPE (Avanti, 850725), Cholesterol (Avanti, 700100), and DMG-PEG2K (Avanti, 880151) at molar ratios of 50:38.5:10:1.5. RNA was incorporated at an N:P ratio of 10. Following encapsulation, the LNPs were dialyzed in sterile 1X Phosphate-buffered saline (PBS) overnight at 4°C. Particle size was determined via dynamic light scattering using a NanoBrook Omni (Brookhaven Instruments). Encapsulation efficiency and RNA concentration were measured with the Quantifluor RNA System (Promega). The LNPs were then concentrated with a 3 kDa MWCO Amicon Ultra Centrifugal filter to approximately 200 ng/µL and further diluted to 125ng/µL with sterile PBS. LNPs were either stored at 4°C and administered within 3 days of preparation or stored at-20°C until use.

### Animal experiment

All animal procedures were approved by the Boston University Institutional Animal Care and Use Committee (IACUC) under protocol PROTO201800680. Both male and female adult C57BL/6 mice (Jackson Labs or Charles River Laboratory) were used, and their ages were 2-5 months at the start of the experiment. During aseptic stereotaxic surgery, mice were anesthetized with isoflurane (5% for induction, 2% or less for maintenance) and preoperatively injected with sustained release buprenorphine (Ethiqa XR).

LNPs loaded with saRNA or mRNA (625 ng total, 5µL at 125 ng/µL) were infused into the striatum through a small craniotomy at 3 depths (ML: +1.8mm, AP: +0.5mm, and 2µL at-2.8mm, 2µL at-2.5mm, and 1µL at-2.2mm), at a rate of 100-250 nL/min via a 10µL syringe (Hamilton Company). At the completion of all 3 infusions, the Hamilton syringe was kept in place for an additional 10 minutes before being retracted. Vetbond tissue adhesive (3M Inc) was then applied to close the incision.

### Histology and immunostaining

Mice were transcardially perfused with 4% paraformaldehyde (PFA) buffered in PBS. Brains were collected, post-fixed in 4% PFA-PBS overnight at 4 °C, and transferred to a solution with 30% sucrose in PBS at 4 °C until sinking. Brains were then sliced coronally at 50µm thickness using a freezing microtome (Leica, SM2010R) or a cryostat (Leica, CM3050S). To assess the efficiency and the time course of saRNA-LNP or mRNA-LNP mediated gene expression, we performed whole-slice imaging using an Olympus VS120 slide scanner microscope at 20X magnification to visualize mCherry or GFP fluorescence.

To quantify neuronal versus astrocyte targeting, we selected 2-3 sections positive for mCherry from each mouse at each desired time point using a fluorescence microscope (Leica, DFC365 FX). The selected tissue sections were then incubated in sodium citrate buffer (10mM, 6.0 pH; Sigma Aldrich, W302600) at 37 °C for 1 hour, and then transferred into a blocking solution (5% goat serum and 0.5% TritonX-100 in PBS) to prevent non-specific protein binding for 1 hour.

Sections were then incubated with primary antibodies against neuron-specific nuclear protein, NeuN (rabbit, 1:500; Abcam, ab177487) and glial fibrillary acidic protein, GFAP (chicken, 1:1000; Novus Biologicals, NBP1-05198) at 4 °C overnight followed by secondary antibodies Alexafluor 488 goat anti-rabbit (1:500; Invitrogen, A11008) and Alexafluor 647 goat anti-chicken (1:500; Invitrogen, A21449) for 2 hours at room temperature. Slices were then mounted using Vectashield antifade mounting medium with DAPI (Vector Laboratories, H-1200). Immunofluorescence was then acquired using an Olympus FV3000 confocal microscope at 40X magnification and 2x zoom.

### Expression mediated by different formulations of LNPs

Three experimenters examined a series of brain sections around the injection site in each animal, and selected one section per animal showing the highest fluorophore expression. For each brain section, we manually segmented the fluorophore-positive region (GFP and mCherry) and the full brain area using the polygon selection tool (ImageJ), and then computed their ratio.

### saRNA-LNP and mRNA-LNP mediated expression area and volume analysis

To estimate the mCherry positive area in each brain section, two experimenters, blind to experimental conditions, independently segmented the areas positive for mCherry in MATLAB. As the segment areas in each slice were largely identical between the two experimenters, the mean area segmented by the two experiments was used for each slice. For each region and timepoint, the two largest mCherry positive areas per animal were compared between the saRNA and mRNA groups using a linear mixed-effects model (Area ∼ Group + (1 | Mouse), with a random intercept per animal to account for the two sections contributed by each mouse. The fixed-effect estimate for group gave the saRNA-versus-mRNA difference and its P value, and P < 0.05 was considered significant.

To estimate saRNA-LNP and mRNA-LNP mediated mCherry expression volume in each mouse, we first registered brain sections with mCherry fluorescence to the Allen Mouse Brain Common Coordinate Framework v3 (25-µm resolution) using DeepSlice, an automated machine-learning method for estimating section position and orientation. Each registration was inspected and refined in QuickNII against the STPT average 2015 reference image. Anterior–posterior (AP) positions (relative to bregma) were calculated from the center of each registered plane.

Manually segmented fluorophore positive areas in the subcortical and corpus callosum regions were then matched to their corresponding registered sections. To estimate the expression volume around the injection site in each mouse, we computed the volume labelled between AP −0.607 and AP +0.254 mm, about 0.25mm posterior to the injection track at AP +0.5 mm, thereby avoiding injection-associated tissue damage that introduce expression variability. Expression area at the boundaries were linearly interpolated between flanking sections, and the volume was estimated by trapezoidal integration of area across sections within the examined AP positions. Mice were grouped into ≤1-week, 2–3-week, and 4–5-week intervals, and saRNA-LNP and mRNA-LNP mediated expression volumes were compared using two-sided Mann-Whitney U tests, with each mouse treated as one independent biological replicate.

### Colocalization analysis

Colocalization analysis was performed using open-source software CellProfiler (https://cellprofiler.org/). Specifically, TIFF images were loaded in CellProfiler as single-channel grayscale images, which were manually thresholded to generate masks corresponding to NeuN positive neurons, GFAP+ glia, or mCherry+ positive objects. As neurons have defined morphology, we applied an additional filter that was optimized by manually adjusting the object area and formfactors to capture the standard shapes of neurons across all NeuN+ neurons. False positive pixels were defined as those positive for more than one marker (NeuN, GFAP, or mCherry) and were removed from all subsequent calculations. To compute the labeling specificity of saRNA in astrocytes, we computed the ratio of pixels positive for both mCherry and GFAP over pixels positive for mCherry. To compute the labeling specificity of saRNA neurons, we computed the ratio of pixels positive for both mCherry and NeuN over the pixels positive for mCherry. Wilcoxon rank-sum tests were used to compare the ratio of neuron and astrocyte labeled since data is not normally distributed and nonparametric.

### Quantification of astrocyte reactivity

GFAP immunofluorescence in each brain section, a measure of astrocyte reactivity, was quantified in Fiji/ImageJ. To identify tissue outline, we applied Gaussian blurring and auto-thresholding of the images, filled small texture holes, and excluded internal cavities (e.g., ventricular space), and obtained the tissue outline as the largest connected object. A midline was then manually drawn to obtain the two hemispheres, ipsilateral and contralateral to the injection side. To identify GFAP positive area, GFAP fluorescence images were background subtracted, filtered with Tubeness filter, and manually thresholded for each brain section. The ratio of GFAP positive area was then computed as that on injected hemisphere over the contralateral hemisphere. Because GFAP positive ratios multiplicative and skewed, analyses were performed on log-transformed ratios, and then back-transformed for reporting as geometric means and fold-changes. We assessed 2–4 sections per mouse in 3–4 mice per group. Differences were assessed a linear mixed-effects model, log(ratio) ∼ group + (1 | mouse), with mouse as a random intercept (MATLAB, fitlme). Each group’s difference from a ratio of 1 was tested using two-sided Wilcoxon signed-rank test, followed by Benjamini-Hochberg post hoc test.

### Human cortical tissue experiment

Human cortical tissue was obtained from surgical resections from epilepsy patients at Boston Children’s Hospital, with informed consent (IRB 09-02-0043), in collaboration with the Repository Core for Neurological Disorders within the Rosamund Stone Zander and Hansjoerg Wyss Translational Neuroscience Center, Boston Children’s Hospital, which is also supported by the IDDRC (NIH P50HD105351). Samples used in this study were collected from macroscopically normal non-lesional cortex.

Immediately following resection, samples were directly placed in carbogenated ice-cold slicing artificial cerebral spinal fluid (ACSF) (80mM NaCl, 75 mM Sucrose, 2.5mM KCl, 1.25mM NaH_2_PO_4_, 26mM NaHCO_3_, 10 mM D-glucose, 0.5mM CaCl_2_, 7mM MgCl_2_, 1.3mM sodium ascorbate, 3.0 mM sodium pyruvate, 1x Penicillin-Streptomycin (Thermo Fisher, 15140122), and 3U/mL Nystatin (Sigma-Aldrich, N1638)) and transported to lab under 15 min. 300 µm thick slices were prepared using a Leica VT1200S vibratome and transferred into a recovery solution (HBSS supplemented with 20mM HEPES, 1 x Penicillin-Streptomycin, and 3U/mL Nystatin, at pH 7.3) at room temperature saturated with 95% O2 5% CO2. After 30min recovery at room temperature, slices were transferred onto Millicell 6-well culture inserts (Millipore PICM0RG50) in culture media (96% (v/v) BrainPhys (StemCell Technologies, 5790), 2.6 mM calcium chloride, 2.6 mM magnesium chloride, 1x N-2 Supplement (Thermo Fisher, 17502001), 1x B-27™ Supplement serum-free (Thermo Fisher, 17504044), 1x Penicillin-Streptomycin (Thermo Fisher, 15140122) 3U/mL Nystatin (Sigma-Aldrich, N1638), 2 μM ascorbic acid, 10 mM D-Glucose, 20ng/mL human recombinant BDNF (StemCell Technologies, 78005), 20ng/mL human recombinant GDNF (StemCell Technologies, 78058) and 20 mM HEPES) in a 6 well plate, and placed in the incubator (5% CO2, 37°C) for an hour before changing to culture media without HEPES and then placed in the incubator overnight. Slices were transduced next day (DIV1) with 4mL of LNPs loaded with saRNA or mRNA by directly applying onto the surface of the slice. Culture media was exchanged every 2-3 days. Z-stack images of live brain slices were acquired with an LSM 980 Airyscan 2 Confocal (Zeiss).

### Human Tissue Expression Quantification

The fraction of mCherry positive area was quantified in Fiji/ImageJ. Briefly, we first subtracted background, and then identified the total tissue area (either the entire field of view, or manually segmented if the tissue did not fill the field of view). mCherry positive pixels were identified using an intensity threshold manually set for each image, as signal and background intensity varied substantially across acquisitions and specimens. For each image we computed the fraction of mCherry positive area over the entire tissue area.

## ACKNOWLEDGEMENTS

X. H. acknowledges funding from NIH 1RF1NS129520, 1R01NS115797, and NSF 2002971-DIOS. M.W.G. acknowledges funding from National Science Foundation TRAILBLAZER; EFMA-2421692, Boston University Kilachand Fund, and William Fairfield Warren Professorship at Boston University. W.W.W. acknowledges funding from the Boston University Kilachand Fund. D.C.S. acknowledges funding from the American Epilepsy Society Predoctoral Fellowship. J.F. acknowledges funding from NIH T32-NS136080. E.S. acknowledges funding from NIH T32-GM008764 and NIH F31 NS143166-01. J.E.M. acknowledges funding from the NSF Graduate Research Fellowship Program 2234657 and NIH T32 Translational Research in Biomaterials Training Program (T32EB006359). E. K. O. acknowledges support from pilot grant funding from the Rosamund Stone Zander and Hansjoerg Wyss Translational Neuroscience Center, Boston Children’s Hospital. M. H. acknowledges funding from the Developmental Neurology Institutional Training Grant, Boston Children’s Hospital, T32 NS007473. Human tissue collection was in collaboration with the Repository Core for Neurological Disorders, Boston Children’s Hospital, supported IDDRC (NIH P50HD105351). This work was additionally supported by Boston University Micro and Nano Imaging Facility through NIH S10OD024993.

## DATA AVAILABILITY

All data and code needed to evaluate and reproduce the results in the paper are present in the paper and/or the Supplementary Materials.

## AUTHOR CONTRIBUTION

Conceptualization: WW, MWG, XH

Major data collection & analysis: JF, JEM, MH

Support for data collection and analysis: EH, SP, DS, YZ, YW, CP, LD, ESA, ZY, KL, SS, HL

Writing – Original Draft: JF, JEM, MH, JF, EKO, WW, MWG, XH

Writing – Review & Editing: All authors

Supervision: WW, MWG, XH

## CONFLICT OF INTEREST

JEM, WW, and MWG hold an equity position in Keylicon Biosciences, a start-up that has licensed the modified saRNA technology from Boston University.

---

**Figure S1.**
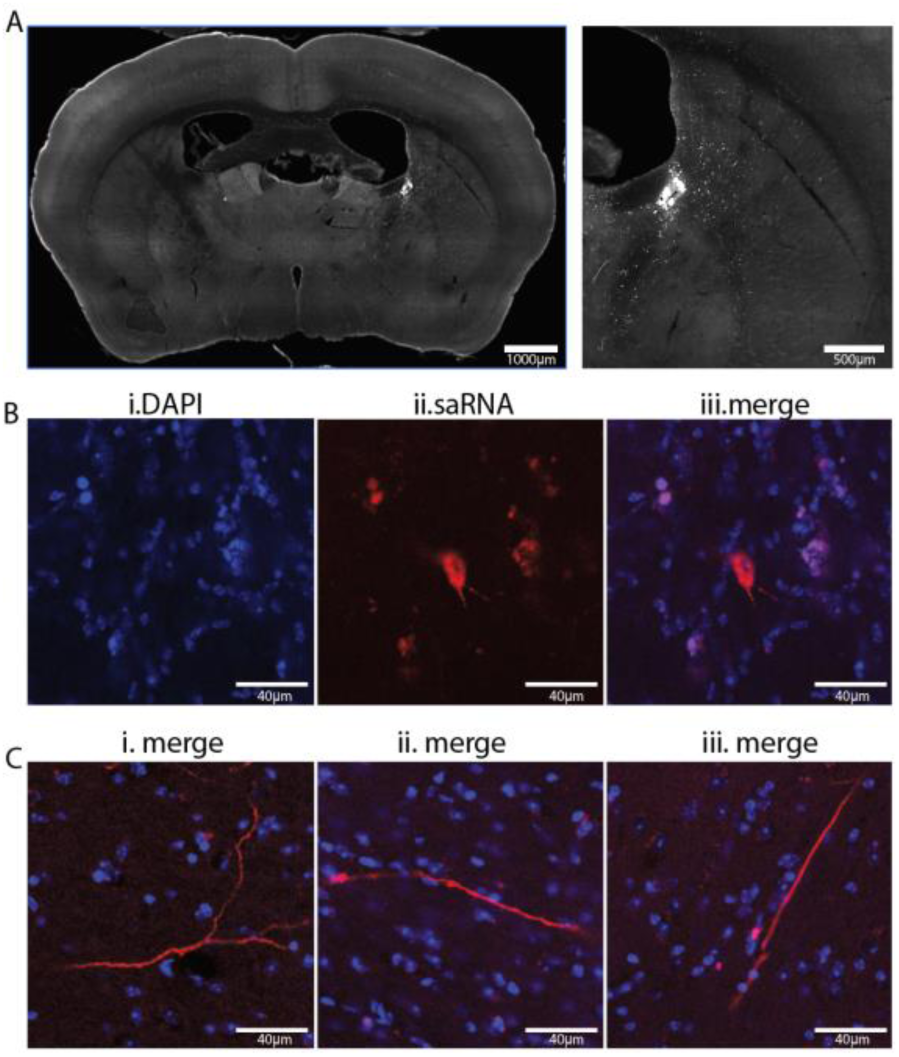
Persistent saRNA-LNP mediated expression in neurons at three months after intra-striatal injection. (A) Coronal section showing the injected hemisphere (left; scale bar, 1000 µm) and a magnified view of the injection site with sparse but bright mCherry+ cells (right; scale bar, 500 µm). (B) Example cell bodies labelled with (i) DAPI, and express (ii) mCherry, and (iii) merge (scale bars, 40 µm). (C) Three additional example images of mCherry+ neuronal processes (red), and DAPI labelled cell bodies (blue) (scale bars, 40 µm).

**Figure. S2.**
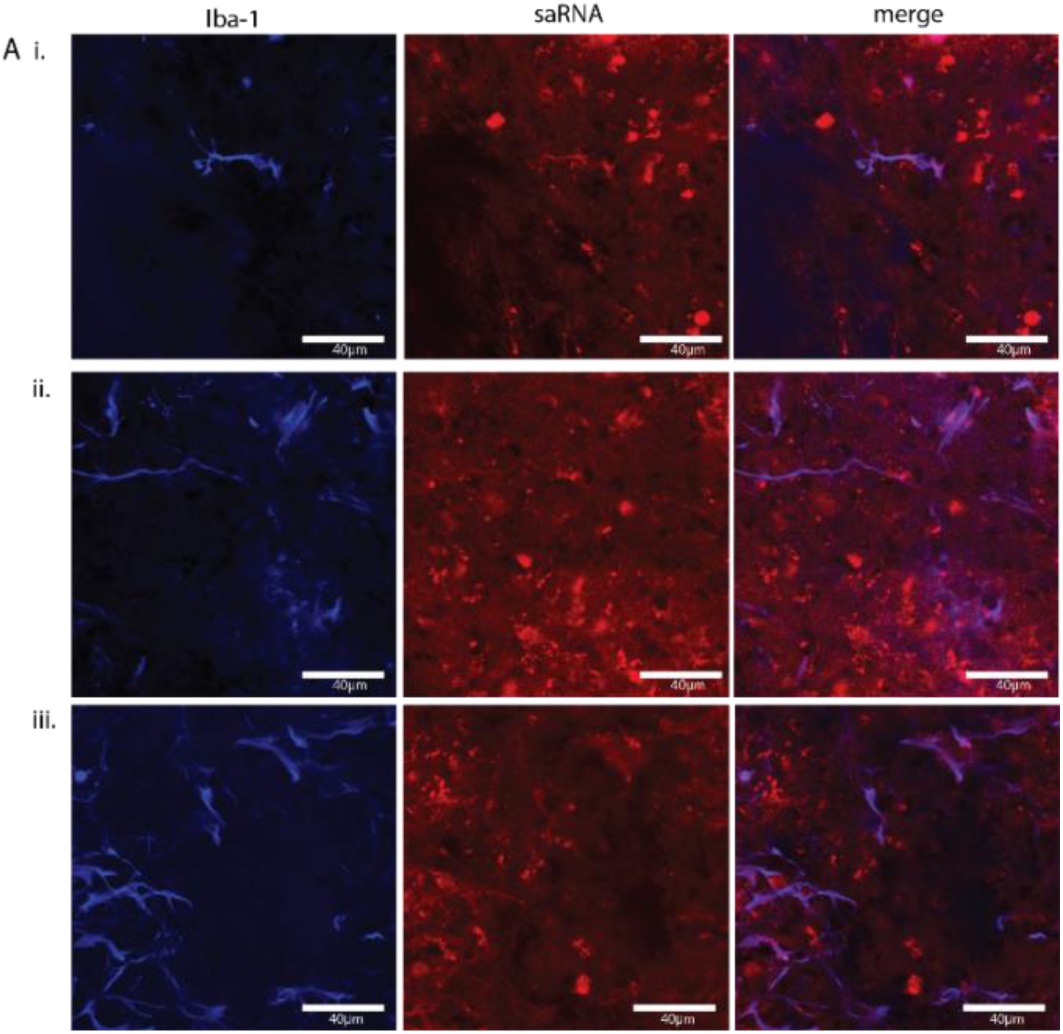
hm5C saRNA-LNPs label microglia around the striatal injection site. Three example images showing Iba-1 (left column), saRNA-mediated mCherry expression (middle column), and the merge (right column). A subset of mCherry+ cells are ramified microglia (scale bars, 40 µm).

**Figure S3.**
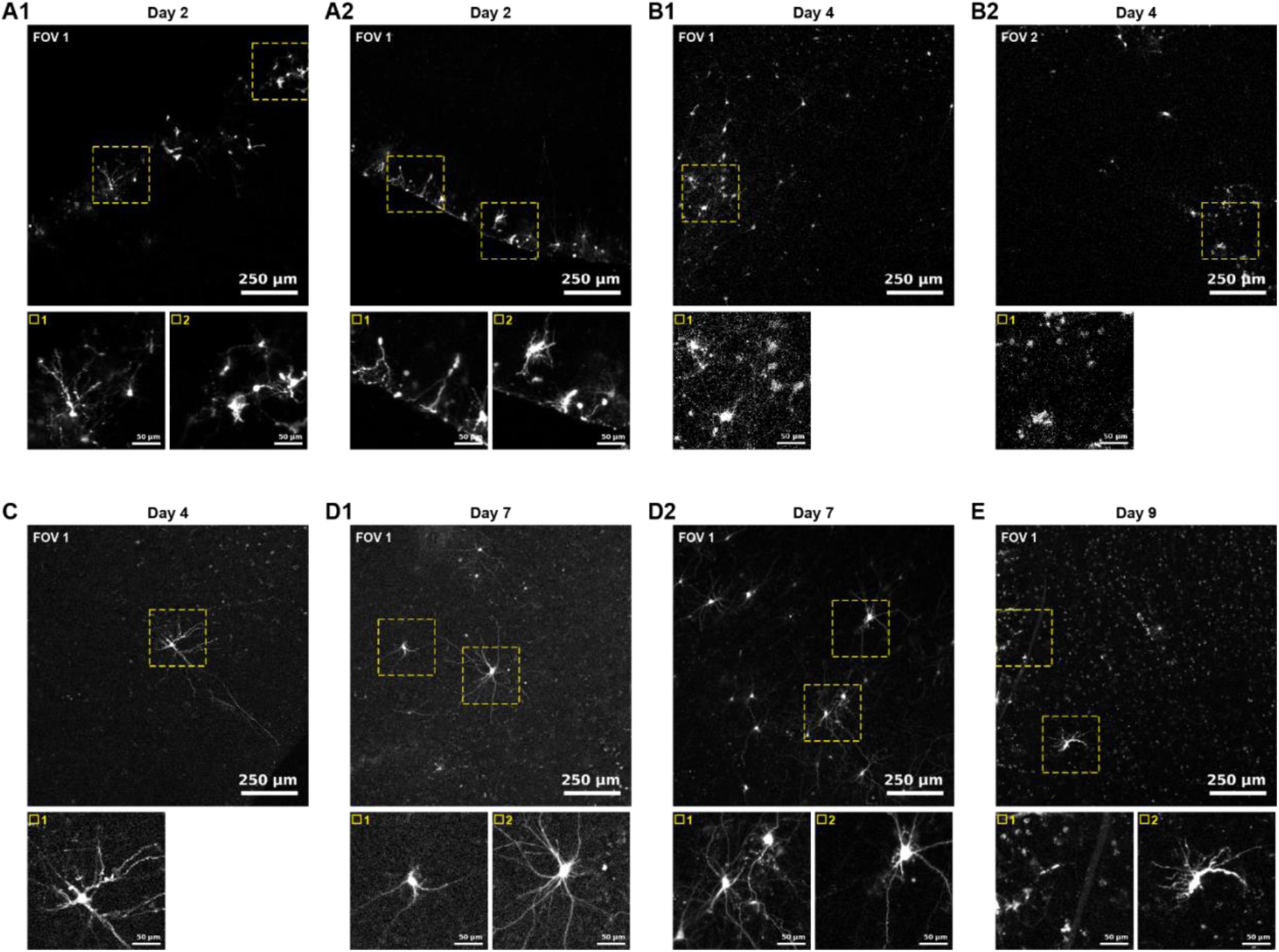
saRNA-LNP mediated mCherry expression in ex vivo human brain slices from five additional patients. (**A-E**), Example mCherry expression in ex vivo slices obtained from 5 addition patients (patient #2 to #6). Top, example fields of view (scale bar, 250 µm). Bottom, zoomed in view of the yellow dashed boxes shown above (scale bars, 50 µm). **(A).** Slice from Patient 2, a 3-year-old male patient (left frontal lobe). (A1) 4 µl dose, 2 days post application. (A2) 4 µl, 2 days post application. **(B)** slice from Patient 3, a 6-month-old female patient (right temporal lobe). 4 µl dose, 4 days post application. (B1, B2) same slice, but two different field of view (FOV1, and FOV2). **(C).** Slice from Patient 4, an 8-year-old female patient (left occipital lobe). 2 µl dose, 4 days post application. (**D).** Slice obtained from Patient 5, a 14-month-old male patient (right temporal lobe). (D1) 2 µl dose, 7 days post application. (D2) 4 µl dose, 7 days post application. **(E).** slice from Patient 6, a 21-year-old female patient (right superior temporal gyrus). 4 µl dose, 9 days post application.

**Figure S4.**
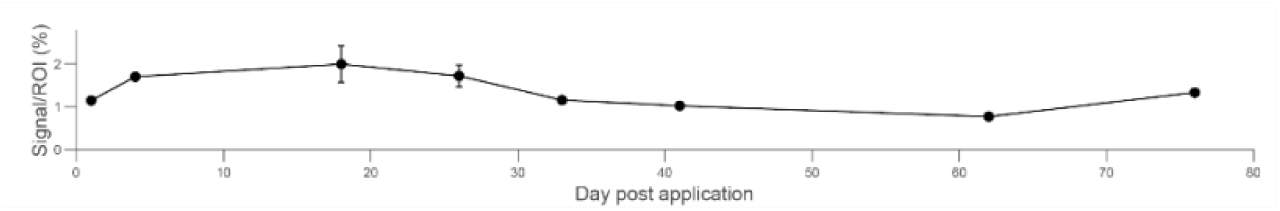
saRNA-mediated transgene expression levels over time in an example ex vivo human brain slice obtained from Patient 1. Expression levels, calculated as the fraction of Cherry+ pixels across the imaging field of view, at varying times following saRNA-LNP application. At day 1, 4, 18, and 26 after application, two fields of view were obtained, and the data shown are mean±standard deviation. All other time points had a single field of view.

**Figure S5.**
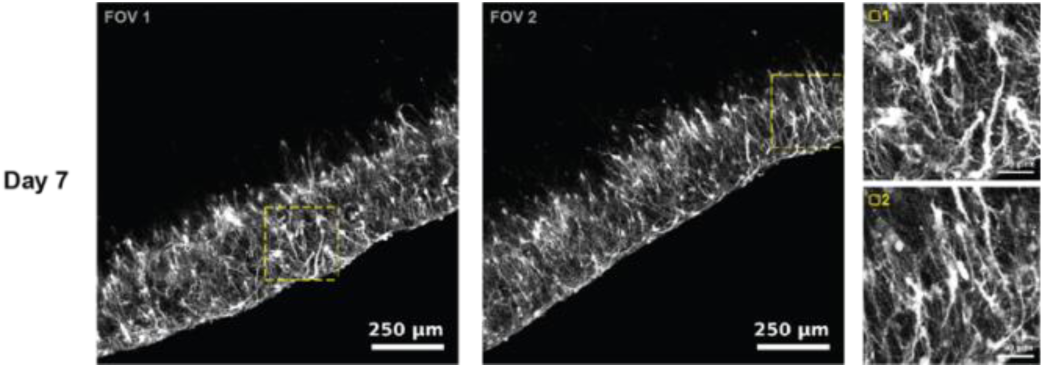
mRNA-LNP mediate transgene expression in an example ex vivo human brain slice. Example images showing mRNA-LNP (4 µl dose) mediated mCherry expression in Patient 3 at 7 days post application. Left two images: example fields of view (FOV1 and FOV2) scale bars, 250 µm. Right, zoomed in view of the yellow dashed boxes shown in FOV1 (top) and FOV 2 (bottom) (scale bars, 50 µm).

**Table S1.** Statistical comparison of saRNA-LNP versus mRNA-LNP mediated mCherry expression volume (related to. **Figure 2 C,D).** n = number of mice per group; two-sided Mann–Whitney rank-sum test. **P < 0.01.

| Region | Time points | Mice number |  | Mann–Whitney |  |
| --- | --- | --- | --- | --- | --- |
|  |  | saRNA | mRNA | U | P values |
| Subcortical | $\leq 1$ week | 5 | 6 | 23 | 0.177 |
| Subcortical | 2–3 weeks | 8 | 8 | 28 | 0.721 |
| Subcortical | 4–5 weeks | 7 | 6 | 42 | 0.0012** |
| Corpus callosum | $\leq 1$ week | 5 | 6 | 17 | 0.792 |
| Corpus callosum | 2–3 weeks | 8 | 8 | 26 | 0.574 |
| Corpus callosum | 4–5 weeks | 7 | 6 | 39 | 0.0082** |

**Table S2.**
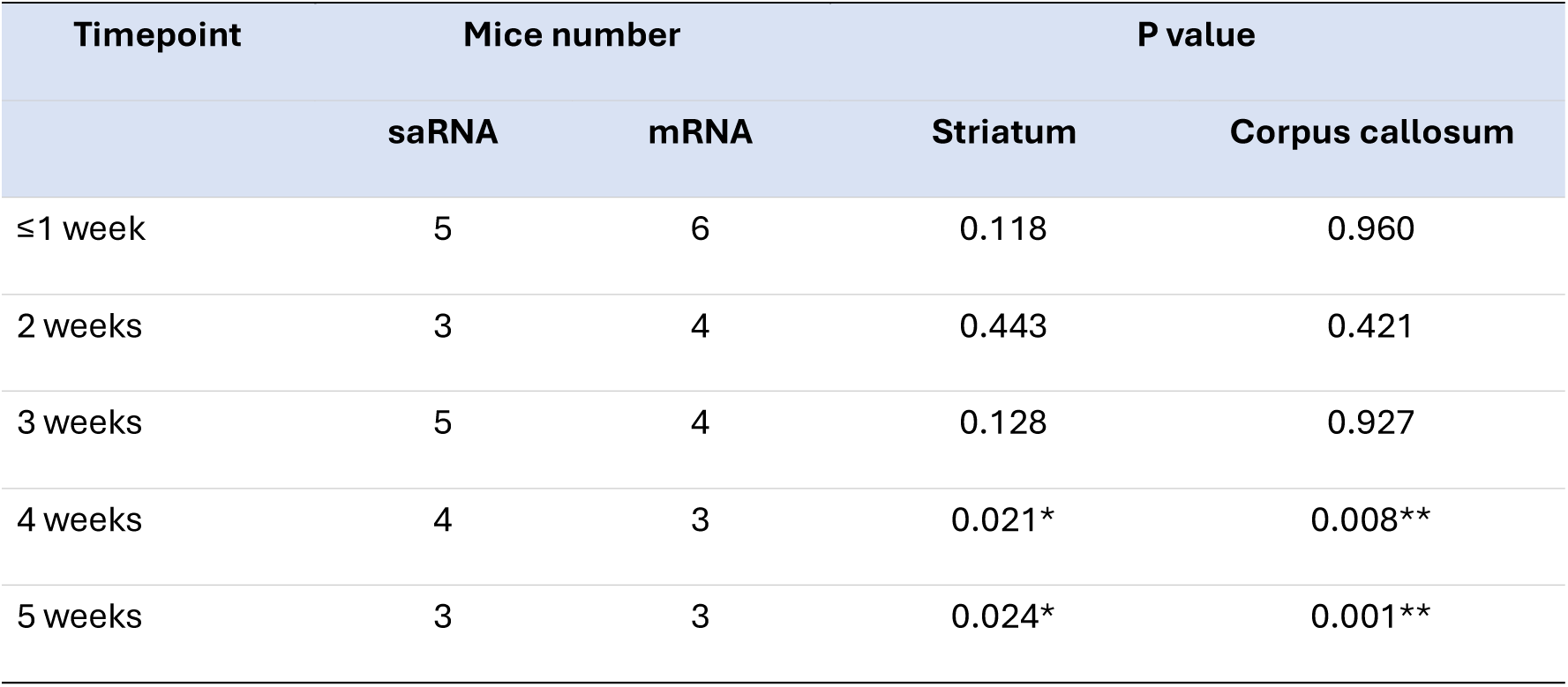
Statistical comparison of saRNA-versus mRNA-mediated expression area (related to. **Fig. 2 E,F).** 2 slices per mouse were included. N=number of mice per group; linear mixed-effects model. *P < 0.05, **P < 0.01.

**Table S3.** Statistical comparison of GFAP+ and NeuN+ labeling (related to. **Fig. 3D).** n=fields of view (FOVs); two-sided Wilcoxon rank-sum test.

| Timepoint | N (FOVs) | P |
| --- | --- | --- |
| 3 days | 19 | $5.4 \times 10^{-5}$ |
| 1 week | 13 | $1.1 \times 10^{-4}$ |
| 2 weeks | 12 | $4.2 \times 10^{-6}$ |
| 3 weeks | 14 | $1.9 \times 10^{-5}$ |
| 4 weeks | 10 | $4.4 \times 10^{-4}$ |

**Table S4.** Statistical comparison of GFAP immunoreactivity across conditions (related to. **Fig. 5C). (a)** GFAP labeling in the injected hemisphere versus the contralateral un-injected hemisphere. N=brain sections; Wilcoxon signed-rank test, followed by by Benjamini–Hochberg post-hoc correction. (B) Comparisons between mice treated with PBS, mRNA-LNP and saRNA-LNP. N=mice numbers; linear mixed-effects model; *P < 0.05, **P < 0.01, ns = not significant.

| *(a) Each group vs contralateral baseline* |  |  | *(b) Pairwise between groups* |  |  |
| --- | --- | --- | --- | --- | --- |
| Group | N sections | P | Comparison (A vs B) | N mice (A vs B) | P |
| PBS | 10 | 0.0029** | PBS vs mRNA | 3 vs 3 | <0.0001** |
| mRNA | 12 | 0.0015** | PBS vs saRNA | 3 vs 4 | <0.0001** |
| saRNA | 9 | 0.0039** | mRNA vs saRNA | 3 vs 4 | 0.866 ns |

## Movie legends

**Movie S1. 3D Illustration of saRNA-mediated mCherry expression.** 3D rendering of the manually segmented mCherry+ expression area across brain sections obtained from a mouse injected with saRNA-LNPs, projected in the reference atlas.

**Movie S2. Illustration of transgene expression volume estimation,** showing manual segmentation of mCherry+ expression area in the subcortical area and the corpus collosum across individual brain sections and their relative locations to the injection site projected in the reference atlas.

## Notes

### Summary of Updates

We have performed more experiments, and the manuscript has been updated to reflect the addition of new results.

